# Association mapping of inflammatory bowel disease loci to single variant resolution

**DOI:** 10.1101/028688

**Authors:** Hailiang Huang, Ming Fang, Luke Jostins, Maša Umićević Mirkov, Gabrielle Boucher, Carl A Anderson, Vibeke Andersen, Isabelle Cleynen, Adrian Cortes, François Crins, Mauro D’Amato, Valérie Deffontaine, Julia Dimitrieva, Elisa Docampo, Mahmoud Elansary, Kyle Kai-How Farh, Andre Franke, Ann-Stephan Gori, Philippe Goyette, Jonas Halfvarson, Talin Haritunians, Jo Knight, Ian C Lawrance, Charlie W Lees, Edouard Louis, Rob Mariman, Theo Meuwissen, Myriam Mni, Yukihide Momozawa, Miles Parkes, Sarah L Spain, Emilie Théâtre, Gosia Trynka, Jack Satsangi, Suzanne van Sommeren, Severine Vermeire, Ramnik J Xavier, International IBD Genetics Consortium, Rinse K Weersma, Richard H Duerr, Christopher G Mathew, John D Rioux, Dermot PB McGovern, Judy H Cho, Michel Georges, Mark J Daly, Jeffrey C Barrett

## Abstract

Inflammatory bowel disease (IBD) is a chronic gastrointestinal inflammatory disorder that affects millions worldwide. Genome-wide association studies (GWAS) have identified 200 IBD-associated loci, but few have been conclusively resolved to specific functional variants. Here we report fine-mapping of 94 IBD loci using high-density genotyping in 67,852 individuals. Of the 139 independent associations identified in these regions, 18 were pinpointed to a single causal variant with >95% certainty, and an additional 27 associations to a single variant with >50% certainty. These 45 variants are significantly enriched for protein-coding changes (n=13), direct disruption of transcription factor binding sites (n=3) and tissue specific epigenetic marks (n=10), with the latter category showing enrichment in specific immune cells among associations stronger in CD and gut mucosa among associations stronger in UC. The results of this study suggest that high-resolution, fine-mapping in large samples can convert many GWAS discoveries into statistically convincing causal variants, providing a powerful substrate for experimental elucidation of disease mechanisms.

---

Inflammatory bowel disease (IBD) is a chronic, debilitating disorder of the gastrointestinal tract with peak onset in adolescence and early adulthood. More than 1.4 million people are affected in the USA alone^1^, with an estimated direct healthcare cost of $6.3 billion/year. IBD affects millions worldwide with a rising prevalence, particularly in pediatric and non-European ancestry populations^2^. IBD is comprised of two etiologically related subtypes, ulcerative colitis (UC) and Crohn’s disease (CD), which have distinct presentations and treatment courses. To date, 200 genomic loci have been associated with IBD^3,4^, but only a handful have been conclusively ascribed to a specific causal variant with direct insight into the underlying disease biology. This scenario is common to all genetically complex diseases, where the pace of identifying associated loci outstrips that of defining specific molecular mechanisms and extracting biological insight from each association.

The widespread correlation structure of the human genome (known as linkage disequilibrium, or LD) often results in similar evidence for association among many nearby variants. However, unless LD is perfect (r^2^ = 1), it is possible, with sufficiently large sample size, to statistically resolve causal variants from neighbors even at high levels of correlation (Extended Data Figure 1 and van de Bunt *et al.*^5^). Novel statistical approaches applied to very large datasets have begun to address this problem^6^ but also require that the highly correlated variants are directly genotyped or imputed with certainty. Truly high-resolution mapping data, when combined with increasingly sophisticated and comprehensive public databases annotating the putative protein-coding and regulatory function of DNA variants, are likely to reveal novel insights into disease pathogenesis^7-9^ and the mechanistic involvement of disease-associated variants.

**Figure 1.**
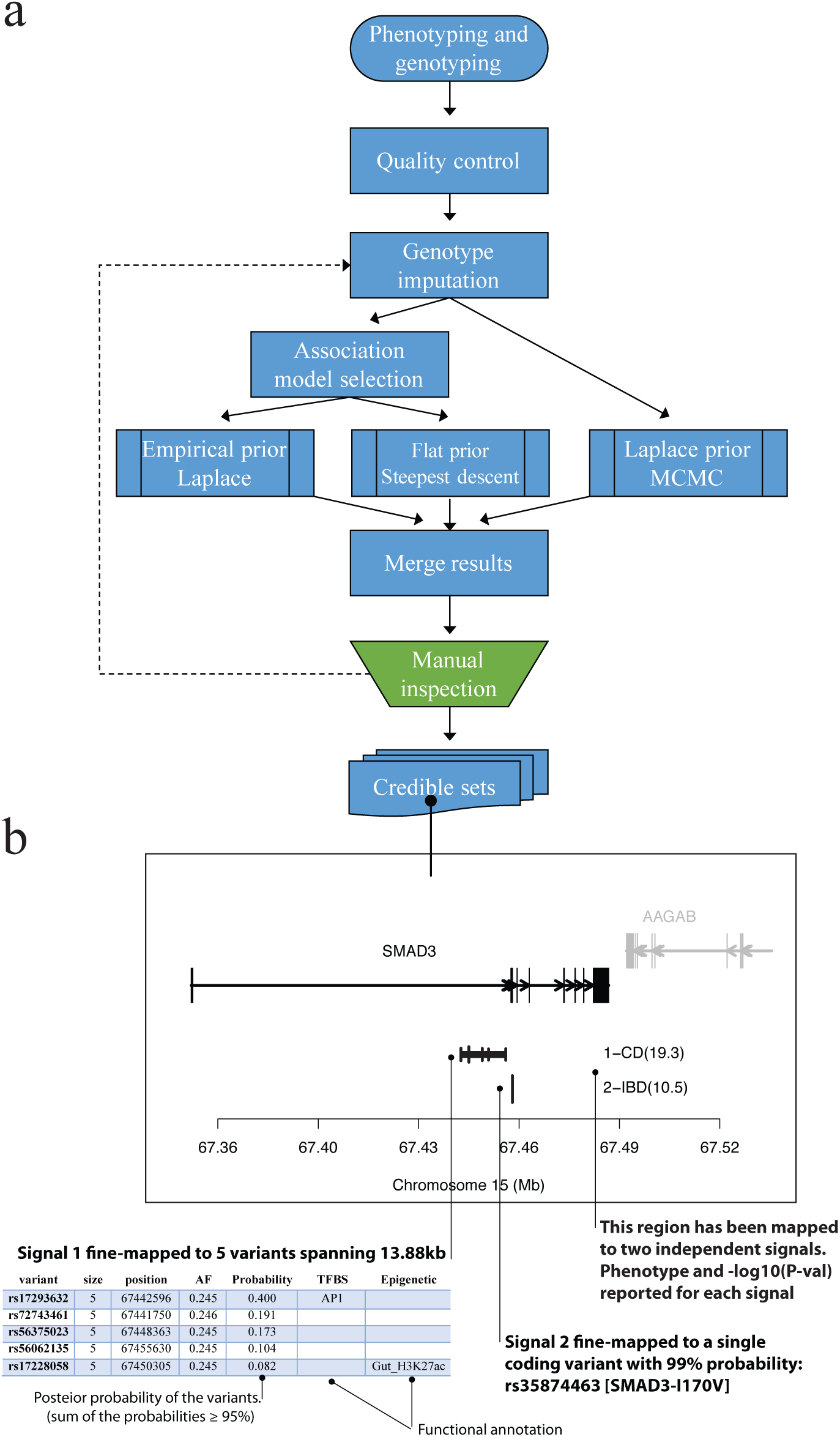
Procedures in the fine-mapping analysis. **a**, Flowchart of fine-mapping steps. Dashed line means the imputation has been performed only once after manual inspection (not iteratively). **b**, An example output from fine-mapping. This region has been mapped to two independent signals. For each signal, fine-mapping reports the phenotype it is associated with, the variants it is fine-mapped to and their posterior probabilities.

## Genetic architecture of IBD associated loci

As part of a large collaborative effort led by the International IBD Genetics Consortium (IIBDGC), 67,852 study subjects of European ancestry, including 33,595 IBD (18,967 CD and 14,628 UC) and 34,257 healthy controls were genotyped using the Illumina™ (San Diego, CA, USA) Immunochip. This custom genotyping array was designed to include all known variants from European individuals in the February 2010 release of the 1000 Genomes Project^10,11^ in 186 high-density regions known to be associated to one or more of 12 immune-mediated diseases^12^. We evaluated ninety-seven of these regions previously associated with IBD^3^ and containing one or more associated variants (p < 10^−6^) in this data set. The major histocompatibility complex was excluded from these analyses as fine-mapping has been reported elsewhere^13^. Because fine-mapping uses subtle differences in strength of association between tightly correlated variants to infer which is most likely to be causal, it is particularly sensitive to data quality. We therefore performed stringent quality control (QC) to remove genotyping errors and batch effects, including manual cluster plot inspection for 905 variants (Methods). After QC, we imputed this dataset using the 1000 Genomes reference panel (December 2013, downloaded from IMPUTE2^14,15^ website) to fill in missing variants or genotype data dropped in chip design or QC (Figure 1a).

We applied three complementary Bayesian fine-mapping methods that used different priors and model selection strategies both to identify independent association signals within a region (Supplementary Methods), and to assign a posterior probability of causality to each variant (Figure 1a). For each independent association signal, we sorted all variants by the posterior probability of association, and added variants to the ‘credible set’ of associated variants until the sum of their posterior probability exceeded 95% - that is, the credible set contains the minimum list of DNA variants that are >95% likely to contain the causal variant (Figure 1b). These sets ranged in size from one to > 400 variants. We merged these results (Methods) and subsequently focused (Figure 1a) only on signals where an overlapping credible set of variants was identified by at least two of the three methods and all variants were either directly genotyped or well imputed (Methods). Fluorescent signal intensity cluster plots were manually reviewed for all variants in credible sets with ten or fewer variants, and a second round of imputation and analysis was performed if any genotypes were removed based on this review.

In 3 out of 97 regions, a consistent credible set could not be identified; when multiple independent effects exist in a region with several highly correlated signals, multiple distinct fine-mapping solutions may not be distinguishable (Supplementary Notes). Sixty-eight of the remaining 94 regions contain a single credible set, while 26 harbored two or more independent association signals, for a total of 139 independent associations defined across the 94 regions (Figure 2a). Only *IL23R* and *NOD2* (both previously established to contain multiple associated protein-coding variants^16^), contain more than three independent signals. Consistent with previous reports^3^, the vast majority of signals are associated with both CD and UC. However, many of these have significantly stronger association with one subtype than the other. For the purposes of enrichment analyses below, we compare 79 signals that are more strongly associated with CD to 23 signals that are more strongly associated with UC (the remaining 37 are equally associated with both subtypes) (“list of credible sets” sheet, Supplementary Table 1).

**Figure 2.**
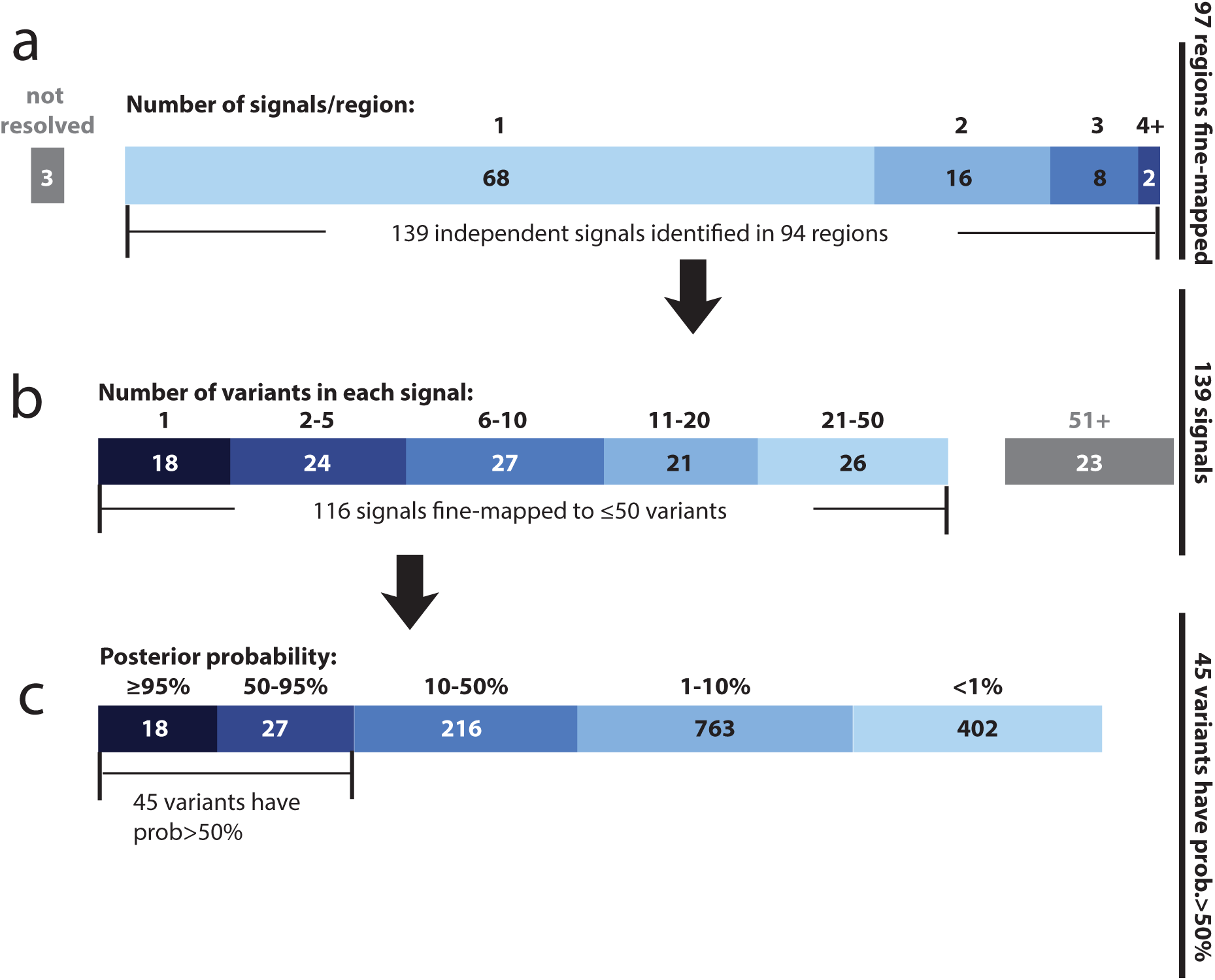
Summary of fine-mapped associations. **a**, sixty-eight loci hosting a single association and 26 loci hosting multiple independent associations. **b**, Number of variants in credible sets. 18 associations were fine-mapped to a single variant, and 116 to ≤ 50 variants. Only credible sets having ≤ 50 variants were advanced for set-enrichment analyses (epigenetics and eQTL). **c**, distribution of the posterior probability in credible sets having ≤ 50 variants. 45 variants have posterior probability > 50% and were advanced for variant-based enrichment analyses (coding, TFBS disruption and epigenetics).

**Table 1.** Summary of variants having posterior probability >50%. Variants were sorted by their posterior probabilities. AF: allele frequency. PROB: posterior probability for being a causal variant. FUNC: functional annotations including coding (C), disrupting transcription factor binding sites (T), Epegenetic peaks (E) and colocalization with eQTL (Q).

| VARIANT | CHR | POSITION | TRAIT | AF | PROB | FUNC | ANNOTATION |
| --- | --- | --- | --- | --- | --- | --- | --- |
| **Signals mapped to a single variant** |  |  |  |  |  |  |  |
| rs7307562 | 12 | 40724960 | CD | 0.398 | 0.999 |  | LRRK2 (intronic) |
| rs2066844 | 16 | 50745926 | CD | 0.063 | 0.999 | C | NOD2(R702W) |
| rs2066845 | 16 | 50756540 | CD | 0.022 | 0.999 | C | NOD2(G908R) |
| rs6017342 | 20 | 43065028 | UC | 0.544 | 0.999 | E | HNF4A (downstream),<br>Gut_H3K27ac |
| rs61839660 | 10 | 6094697 | CD | 0.094 | 0.999 | E | IL2RA (intronic),<br>Immune_H3K4me1 |
| rs5743293 | 16 | 50763781 | CD | 0.964 | 0.999 | C | fs1007insC |
| rs6062496 | 20 | 62329099 | IBD | 0.587 | 0.996 | T | RTEL1-TNFRSF6B (ncRNA_intronic),<br>EBF1 TFBS |
| rs141992399 | 9 | 139259592 | IBD | 0.005 | 0.995 | C | CARD9(1434+1G>C) |
| rs35667974 | 2 | 163124637 | UC | 0.021 | 0.994 | C | IFIH1(I923V) |
| rs74465132 | 7 | 50304782 | IBD | 0.034 | 0.994 | T,E | IKZF1 (upstream),<br>EP300 TFBS,<br>Immune_H3K4me1 |
| rs4676408 | 2 | 241574401 | UC | 0.508 | 0.994 |  | GPR35 (downstream) |
| rs5743271 | 16 | 50744688 | CD | 0.007 | 0.993 | C | NOD2(N289S) |
| rs10748781 | 10 | 101283330 | IBD | 0.55 | 0.990 | E | NKX2-3 (upstream),<br>Gut_H3K27ac |
| rs35874463 | 15 | 67457698 | IBD | 0.054 | 0.989 | C,E | SMAD3(I170V),<br>Gut_H3K27ac |
| rs72796367 | 16 | 50762771 | CD | 0.023 | 0.983 |  | NOD2 (intronic) |
| rs1887428 | 9 | 4984530 | IBD | 0.603 | 0.974 |  | JAK2 (upstream) |
| rs41313262 | 1 | 67705900 | CD | 0.014 | 0.973 | C | IL23R(V362I) |
| rs28701841 | 6 | 106530330 | CD | 0.116 | 0.971 |  | PRDM1 (upstream) |
| **Signals mapped to 2-50 variants but the lead variant have posterior probability > 50%** |  |  |  |  |  |  |  |
| rs76418789 | 1 | 67648596 | CD | 0.006 | 0.937 | C | IL23R(G149R) |
| rs7711427 | 5 | 40414886 | CD | 0.633 | 0.919 |  |  |
| rs1736137 | 21 | 16806695 | CD | 0.407 | 0.879 |  |  |
| rs104895444 | 16 | 50746199 | CD | 0.003 | 0.865 | C | NOD2(V793M) |
| rs56167332 | 5 | 158827769 | IBD | 0.353 | 0.845 |  | IL12B |
| rs104895467 | 16 | 50750810 | CD | 0.002 | 0.833 | C | NOD2(N852S) |
| rs630923 | 11 | 118754353 | CD | 0.153 | 0.820 |  |  |
| rs3812565 | 9 | 139272502 | IBD | 0.402 | 0.815 | Q | eQTL of INPP5E in CD4 and CD8;<br>CARD9 in CD14, SEC16A in CD15 |
| rs4655215 | 1 | 20137714 | UC | 0.763 | 0.784 | E | Gut_H3K27ac |
| rs145530718 | 19 | 10568883 | CD | 0.023 | 0.762 |  |  |
| rs6426833 | 1 | 20171860 | UC | 0.555 | 0.752 |  |  |
| chr20:<br>43258079 | 20 | 43258079 | CD | 0.041 | 0.736 |  |  |
| rs17229679 | 2 | 199560757 | UC | 0.028 | 0.716 |  |  |
| rs4728142 | 7 | 128573967 | UC | 0.448 | 0.664 | E | Immune_H3K4me1 |
| rs2143178 | 22 | 39660829 | IBD | 0.157 | 0.662 | T,E | NFKB TFBS, Gut_H3K27ac |
| rs34536443 | 19 | 10463118 | CD | 0.038 | 0.649 | C | TYK2(P1104A) |
| rs138425259 | 16 | 50663477 | UC | 0.009 | 0.648 |  |  |
| rs146029108 | 9 | 139329966 | CD | 0.036 | 0.643 |  |  |
| rs12722504 | 10 | 6089777 | CD | 0.26 | 0.615 |  |  |
| rs60542850 | 19 | 10488360 | IBD | 0.17 | 0.591 |  |  |
| rs2188962 | 5 | 131770805 | CD | 0.44 | 0.590 | E,Q | Gut_H3K27ac,<br>eQTL of SLC22A5 in CD14, CD15 and<br>IL |
| rs2019262 | 1 | 67679990 | IBD | 0.4 | 0.586 |  |  |
| rs3024493 | 1 | 206943968 | IBD | 0.171 | 0.537 | E | Immune_H3K4me1 |
| rs7915475 | 10 | 64381668 | CD | 0.304 | 0.528 |  |  |
| rs77981966 | 2 | 43777964 | CD | 0.077 | 0.521 |  |  |
| rs9889296 | 17 | 32570547 | CD | 0.264 | 0.512 |  |  |
| rs2476601 | 1 | 114377568 | CD | 0.908 | 0.508 | C | PTPN22(W620R) |

Using a restricted maximum likelihood mixed model approach^17^, we evaluated the proportion of total variance in disease risk attributed to these 94 regions and how much of that is explained by the 139 specific associations. We estimated that 25% of CD risk was explained by the specific associations described here, out of a total of 28% explained by these loci (the corresponding numbers for UC are 17% out of 22%). This indicates that our credible sets capture most of the IBD genetic risk at these loci. The single strongest signals in each region contribute 76% of this variance explained and the remaining associations contribute 24% (Extended Data Figure 2b), highlighting the importance of secondary and tertiary associations in the articulation of GWAS results^13,18^.

## Associations mapped to a single variant

For 18 independent signals, the 95% credible set consisted of a single variant (hereafter referred to as ‘single variant credible sets’) and for 24 others, the credible set consisted of two to five variants (Figure 2b). The single variant credible sets included five previously reported coding variants: three in *NOD2* (fs1007insC, R702W, G908R), a rare protective allele in *IL23R* (V362I) and a splice variant in *CARD9* (c.IVS11+1G>C) ^16,19^. The remaining single variant credible sets were comprised of three missense variants (I170V in *SMAD3*, I923V in *IFIH1* and N289S in *NOD2*), four intronic variants (in *IL2RA, LRRK2, NOD2* and *RTEL1/TNFRSF6B*) and six intergenic variants (located 3.7kb downstream of *GPR35;* 3.9kb upstream of *PRDM1;* within a EP300 binding site 39.9 kb upstream of *IKZF1;* 500 bp before the transcription start site of *JAK2;* 9.4kb upstream of *NKX2-3;* and 3.5kb downstream from *HNF4A*) (Table 1). A customizable browser (https://atgu.shinyapps.io/Finemapping) enabling review of the detailed fine-mapping results in each region along with all annotations discussed below has been prepared. Of note, while physical proximity does not guarantee functional relevance, the credible set of variants for 29 associated loci now resides within 50 kb of only a single gene – improved from only 3 so refined using an earlier HapMap-based definition. Using the same definitions, the total number of potential candidate genes was reduced from 669 to 331. Examples of IBD candidate genes clearly prioritized in our data are described in the Supplementary Box.

## Sequence-level consequences of associated variants – protein coding variation

We first annotated the possible functional consequences of the IBD variants by their effect on the amino acid sequences of proteins. Thirteen out of 45 variants that have >50% posterior probability are non-synonymous (Table 1 and Figure 2c), an 18-fold enrichment (p-value=2x10^−13^, Fisher’s exact test) relative to randomly drawn variants in our regions. By contrast, only one variant with >50% probability is synonymous (p=0.42). All common coding variants previously reported to affect IBD risk are included in a 95% credible set including: *IL23R* (R381Q, V362I and G149R); *CARD9* (c.IVS11+1G>C and S12N); *NOD2* (S431L, R702W, V793M, N852S and G908R, fs1007insC); *ATG16L1* (T300A); *PTPN22* (R620W); and *FUT2* (W154X). While this enrichment of coding variation (Figure 3a) provides assurance about the accuracy of our approach, it does not suggest that 30% of all associations are caused by coding variants; rather, it is almost certainly the case that associated coding variants have stronger effect sizes, making them more amenable to fine mapping.

**Figure 3.**
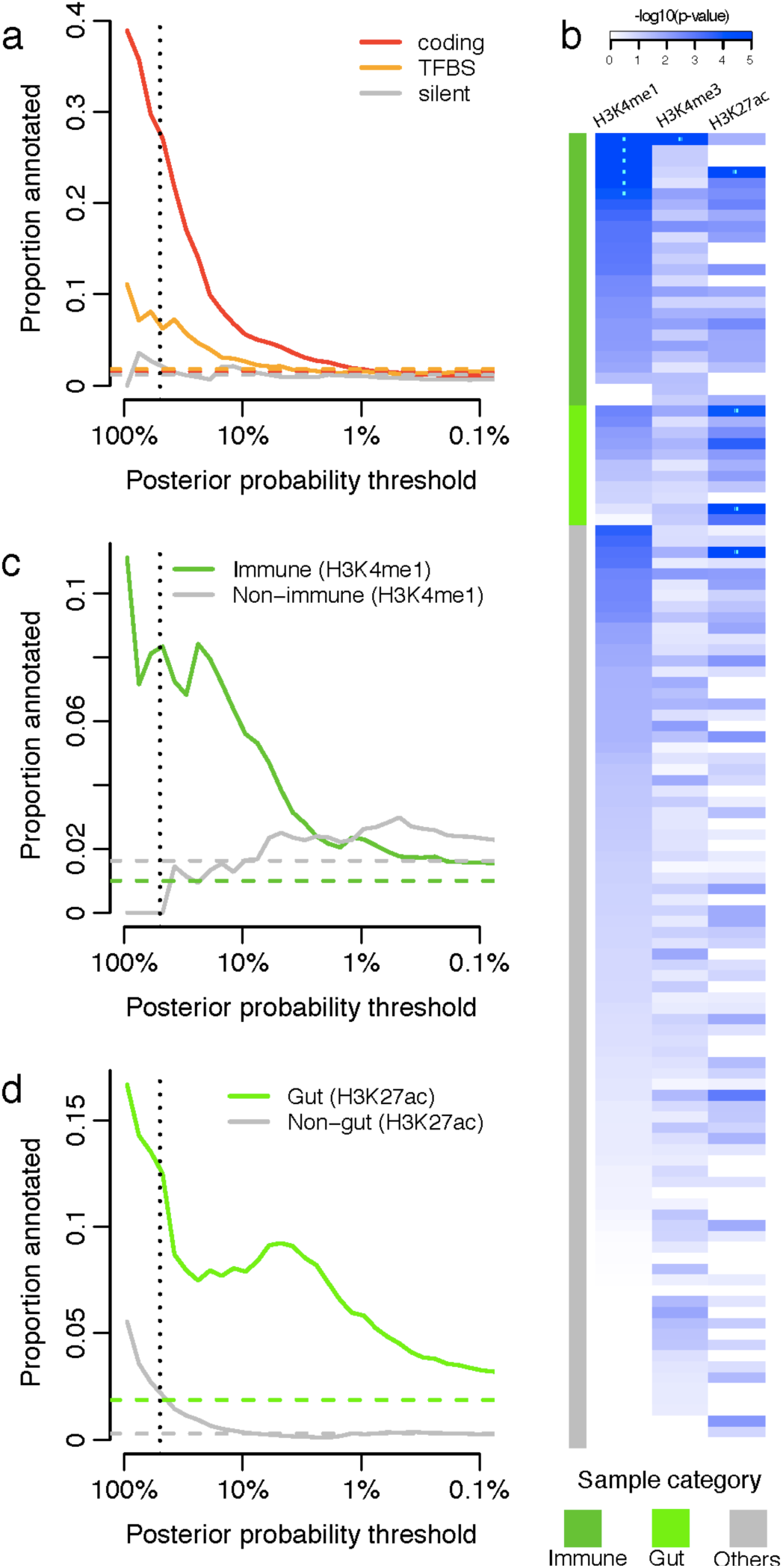
Functional annotation of causal variants. **a**, Proportion of variants that are protein coding, disrupting/creating transcription factor binding motifs (TFBS) or synonymous. **b**, Epigenetic peaks overlapping credible variants in various cell lines. Sample categories were taken from the Roadmap Epigenomics Consortium^36^. Significant cell line-peak pairs have been marked with asterisks. **c**, Proportion of credible variants that overlap H4K4Me1 peaks. **d**, Proportion of credible variants that overlap H3K27ac peaks. In panels **a, c** and **d**, the vertical dotted lines mark the 50% probability and the horizontal dashed lines show the background proportions of each functional category.

## Sequence-level consequences of associated variants – non-coding variation

We next examined the best understood non-coding aspect of DNA sequence: conserved nucleotides in high confidence binding site motifs of 84 transcription factor (TF) families^20^ (Methods). There was a significant positive correlation between TF motif disruption and IBD association posterior probability (p-value=0.006, binomial regression) (Figure 3a), including three variants with >50% probability (two >95%). In the *RTEL1/TNFRSF6B* region, rs6062496 region is predicted to disrupt a TF binding site (TFBS) for EBF1 and overlaps DNaseI hypersensitivity clusters. EBF1 is a TF involved in the maintenance of B cell identity and prevention of alternative fates in committed cells^21^. The second example, rs74465132, is a low frequency (3.6%) protective variant that creates a binding site for EP300 less than 40kbp upstream of *IKZF1* (zinc-finger DNA binding protein). The third notable example of TFBS disruption, although not in a single variant credible set, is detailed in the Supplementary Box for the association at *SMAD3.*

Recent studies have shown that trait associated variants are enriched for epigenetic marks highlighting cell type specific regulatory regions ^22,23^. We compared our credible sets with ChIPseq peaks corresponding to chromatin immunoprecipitation with H3K4me1, H3K4me3 and H3K27ac in 120 adult and fetal tissues, assayed by the NIH Roadmap Epigenomics Mapping Consortium^24^ (Figure 3b). Using a threshold of p=1.3×10^−4^ (0.05 corrected for 360 tests), we observed significant enrichment of H3K4me1 in 6 immune cell types and for H3K27ac in 3 gastrointestinal (GI) samples (sigmoid colon and colonic and rectal mucosa) (Figure 3b and Supplementary Table 2). Furthermore, the subset of signals that are more strongly associated with CD overlap more with immune cell chromatin peaks, whereas UC signals overlap more with GI chromatin peaks (Supplementary Table 2).

These three chromatin marks are correlated both within tissues (we observe additional signal in other marks in the tissues described above) and across related tissues. We therefore defined a set of “core immune peaks” for H3K4me1 and “core GI peaks” for H3K27ac as the set of overlapping peaks in all enriched immune cell and GI tissue types, respectively. These two tracks (immune-K4me1 and gut-K27ac) are independently significant and capture the observed enrichment compared to “control peaks” made up of the same number of ChIPseq peaks across our 94 regions in non-immune and non-GI tissues (Figure 3c,d). These two tracks summarize our epigenetic-GWAS overlap signal, and the combined excess over the baseline suggests that a substantial number of regions, particularly those not mapped to coding variants, may ultimately be explained by functional variation in recognizable enhancer/promoter elements.

## Overlap of IBD credible sets with expression QTLs

Variants that change enhancer or promoter activity might precipitate changes in gene expression, and baseline expression of many genes has been found to be regulated by genetic variation^25-27^. Indeed, these so-called expression quantitative trait loci (eQTLs) have been suggested to underlie a large proportion of GWAS associations ^25,28^. We therefore searched for variants that are both in an IBD associated credible set with 50 or fewer variants and the most significantly associated eQTL variant for a gene in the GODOT study^29^ of peripheral blood mononuclear cells (PBMC) from 2,752 twins. Sixty-eight of the 76 regions with signals fine-mapped to < 50 variants harbor at least one significant eQTL (defined as influencing expression of a gene within 1 Mb of the region with a p-value < 10^−5^). Despite this apparent abundance of eQTLs in fine-mapped regions, only 3 credible sets overlap eQTLs, compared with 3.7 expected by chance (Methods). Data from a more recent independent study (Westra *et al*.)^30^ using PBMCs from 8,086 individuals did not yield a substantively different outcome, demonstrating a modest but non-significant enrichment (8 observed overlaps, 4.2 expected by chance, p=0.07). Using a more lenient definition of overlap which requires the lead eQTL variant to be in LD (R^2^ > 0.4) with an IBD credible set variant increased the number of potential overlaps but again these numbers were not greater than chance expectation (GODOT: observed 14, expected 12.2; Westra *et al.:* observed 11, expected 9.1).

As PBMCs are a heterogeneous collection of immune cell populations, cell type-specific signals, or signals corresponding to genes expressed most prominently in non-immune tissues, may be missed. We therefore tested the enrichment of eQTLs that overlap credible sets in 5 primary T cell populations (CD4+, CD8+, CD19+, CD14+ and CD15+), platelets, and 3 distinct intestinal locations (rectum, colon and ileum) isolated from 350 healthy individuals (ULg dataset, Methods). We observed a significant enrichment of credible SNP/eQTL overlaps in CD4+ cells and ileum (Extended Table 1): 3 and 2 credible sets overlapped eQTLs, respectively, compared to 0.4 and 0.3 expected by chance (p-value=0.007 and 0.025). An enrichment was also observed for the naïve CD14+ cells from another study^31^ (Knight dataset, Extended Data Table 1): eight overlaps observed compared to 2.7 expected by chance (p-value=0.005). We did not observe enrichment of overlaps in stimulated (with interferon or lipopolysaccharide) CD14+ cells from the same source (Extended Data Table 1).

To more deeply investigate eQTL overlaps we applied two colocalization approaches (one based on permutations, one Bayesian, Methods) to eQTL datasets where primary genotype and expression data were available (ULg dataset). We confirmed greater than expected overlap with eQTLs in CD4+ and ileum described above (Figure 4 and Extended Data Table 1). The number of colocalizations in other purified cell types/tissues was largely indistinguishable from what we expect under the null using either method, except for moderate enrichment in rectum (4 observed and 1.4 expected, p=0.039) and colon (3 observed and 0.8 expected, p=0.04). Of these robust colocalizations, only two correspond to an IBD variant with causal probability > 50% (Table 1 and Extended Data Figure 3a).

**Figure 4.**
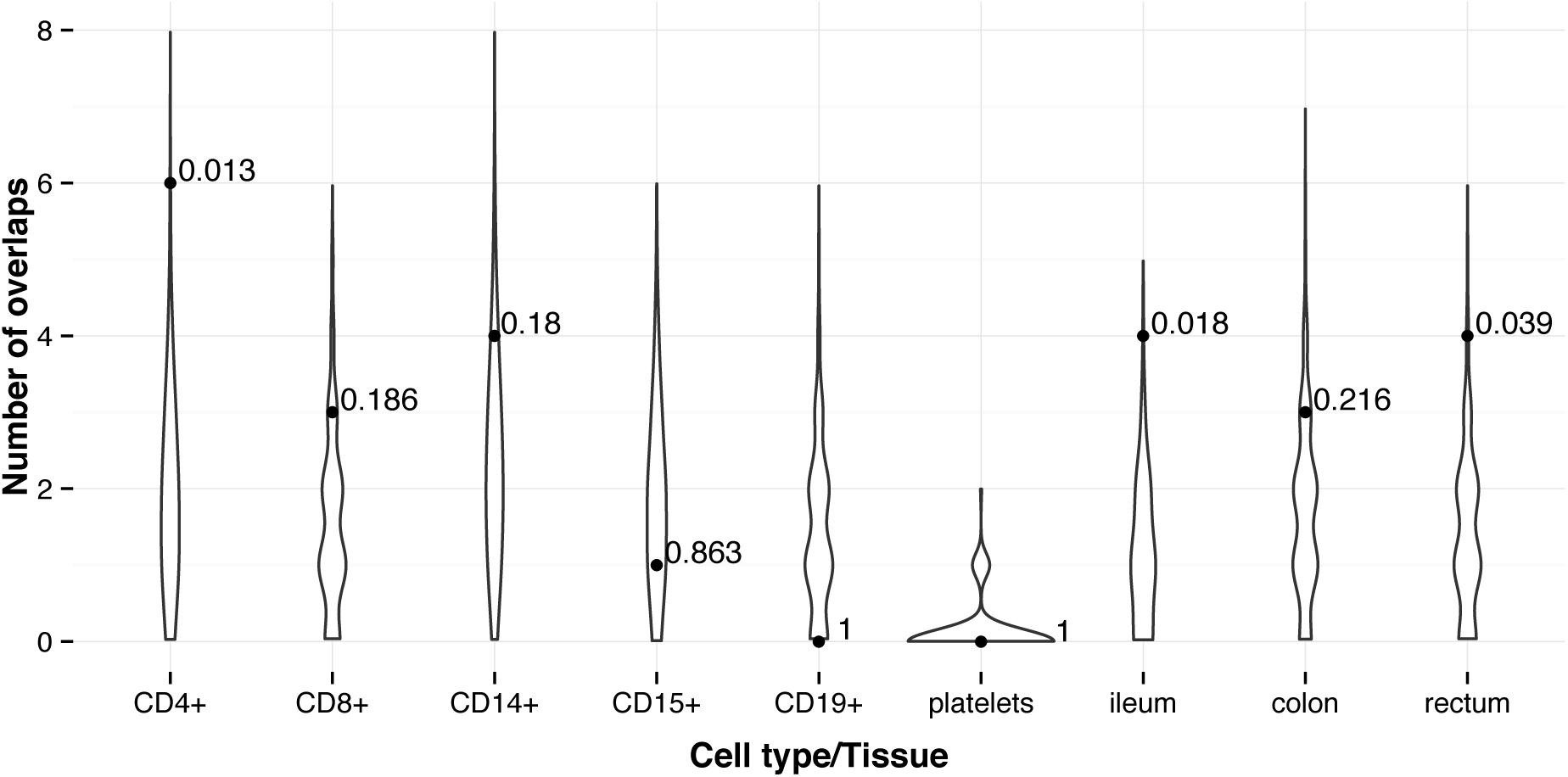
Number of credible sets that colocalize eQTLs. The violin plot shows the distribution of the number of colocalizations by chance (background) and the solid points shows the observed number of colocalizations. P-values of the enrichment were shown next to the solid points. Both the background and the observed numbers were calculated using the permutation based approach (Methods).

## Discussion

We have performed fine-mapping of 94 previously reported genetic risk loci for IBD. Rigorous quality control followed by a integration of three novel fine-mapping methods was employed to generate a list of genetic variants accounting for 139 independent associations across these loci. These associations account for more than 80% of the total variance explained by these loci. Our results substantially improve on previous fine-mapping efforts using a preset LD threshold (e.g. r^2^> 0.6^32^) (Figure 5) by formally modeling the posterior probability of association of every variant. Much of this resolution derives from the very large sample size we employed, because the number of variants in a credible set significantly decreases with increasing test statistics (p-value = 0.0069, Extended Data Figure 4). For example, at 10% allele frequency, 31% of signals are fine-mapped to ≤ 5 variants – this improves to 53% if the sample size were to double again.

**Figure 5.**
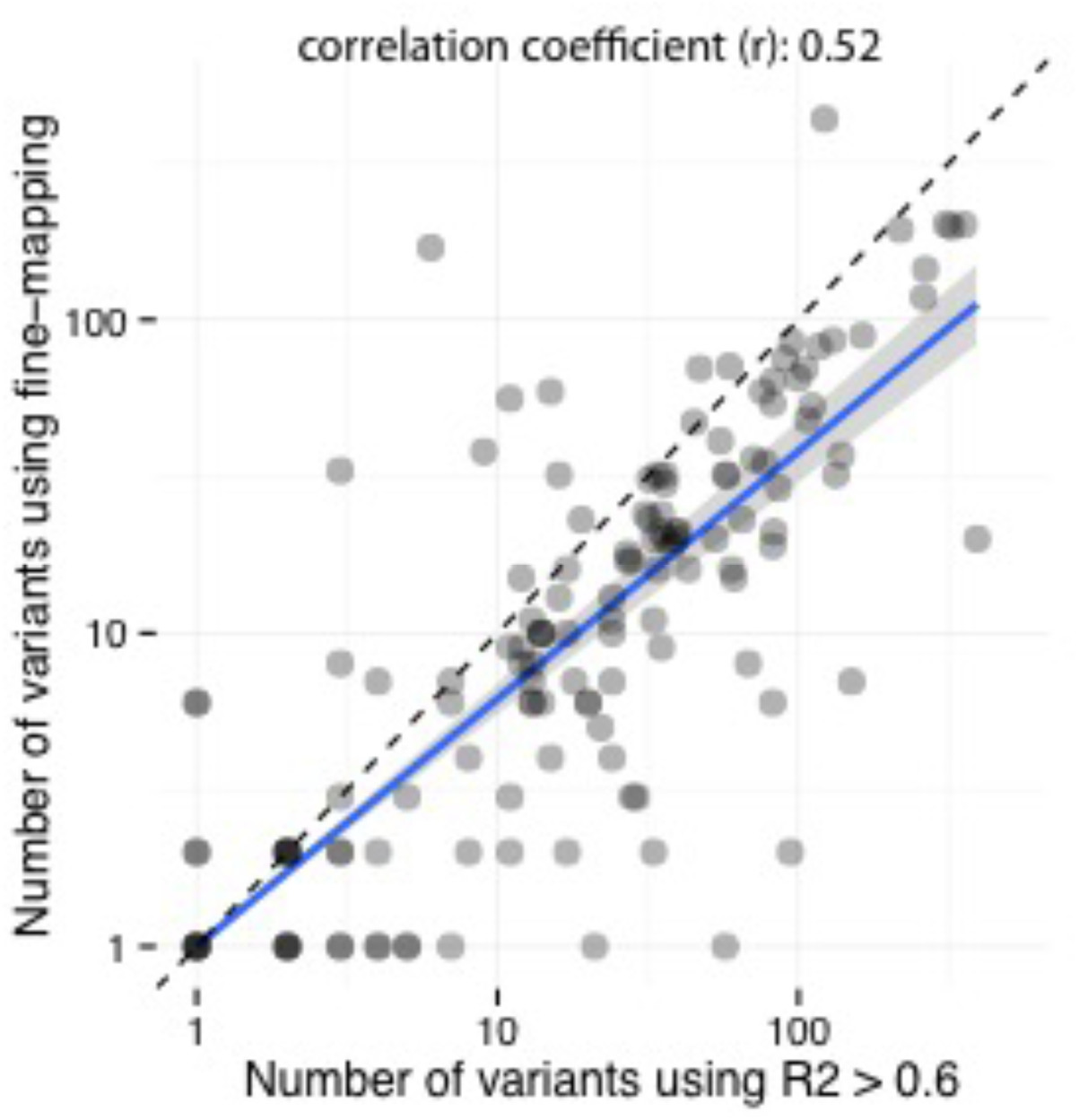
Fine-mapping improved the resolution of genetic associations. We compare the numbers of variants that are mapped in each independent signal using the fine-mapping approach (y axis) and the *R*^2^ > 0.6 cut-off (x axis). Fine-mapping maps most signals to smaller numbers of variants.

Additionally, the high-density of genotyping also aids in improved resolution. For instance, the primary association at IL2RA has now been mapped to a single variant associated with CD, rs61839660. This variant was not present in the Hapmap 3 reference panel and was therefore not reported in earlier studies^3,33^ (nearby tagging variants, rs12722489 and rs12722515, were reported instead). Imputation using the 1000 genomes reference panel and the largest assembled GWAS dataset^3^ did not separate rs61839660 from its neighbors (unpublished results), due to the loss of information in imputation using the limited reference. Only direct genotyping, available in the immunochip high-density regions, permitted the conclusive identification of this as the causal variant.

Accurate fine-mapping should, in many instances, ultimately point to the same variant across diseases in shared loci. Among our single-variant credible sets, we fine-mapped a UC association to a rare missense variant (I923V) in *IFIH1*, which is also associated with type 1 diabetes (T1D)^34^ with an opposite direction of effect (Supplementary Box). The intronic variant noted above (rs61839660, AF=9%) in *IL2RA* was also similarly associated with T1D, again with a discordant directional effect^35^ (Supplementary Box). Simultaneous high-resolution fine-mapping in multiple diseases should therefore better clarify both shared and distinct biology.

High-resolution fine-mapping demonstrates that causal variants are significantly enriched for variants that alter protein coding variants or disrupt transcription factor binding motifs. Enrichment was also observed in H3K4me1 marks in immune related cell types and H3K27ac marks in sigmoid colon and rectal mucosal tissues – with CD loci demonstrating a stronger immune signature and UC loci more enriched for gut tissues. By contrast, overall enrichment of eQTLs is quite modest compared with prior reports and not seen in excess of chance in our well-refined credible sets. This result underscores not only the importance of the high-resolution mapping but also the careful incorporation of the high background rate of eQTLs. It is worth noting that evaluating the overlap between two distinct mapping results is fundamentally different than comparing genetic mapping results to fixed genomic features, and depends on both mappings being well-resolved. While these data strongly challenge the paradigm that easily surveyed baseline eQTLs explain a large proportion of non-coding GWAS signals, the modest excesses observed in smaller but cell-specific data sets suggest that much larger tissue or cell-specific studies (and under the correct stimuli or developmental time points) will resolve the contribution of eQTLs to GWAS hits.

Resolving multiple independent associations may often help target the causal gene more precisely. For example, the *SMAD3* locus hosts a non-synonymous variant and a variant disrupting the conserved transcription factor binding site (also overlapping the H3K27ac marker in gut tissues), unambiguously articulating a role in disease and providing an allelic series for further experimental inquiry. Similarly, the *TYK2* locus has been mapped to a non-synonymous variant and a variant disrupting a conserved transcription factor binding site (Extended Data Figure 5).

One-hundred and sixteen associations have been fine-mapped to ≤ 50 variants. Among them, 27 associations contain coding variants, 20 contain variants disrupting transcription factor binding motifs, and 45 are within histone H3K4me1 or H3K27ac marked DNA regions. However, 40 non-coding associations were not mapped to any known function (Extended Data Figure 3b) despite extensive efforts to integrate with all available annotation, epigenetic and eQTL data.

The best-resolved associations - 45 variants having >50% posterior probabilities for being causal (Table 1) – are similarly significantly enriched for variants with known or presumed function from genome annotation. Of these, 13 variants cause non-synonymous change in amino acids, 3 disrupt a conserved TF binding motif, 10 are within histone H3K4me1 or H3K27ac marked DNA regions in disease-relevant tissues, and 2 co-localize with a significant cis-eQTL (Extended Data Figure 3a).

This analysis leaves, however, 21 non-coding variants, all of which have extremely high probabilities to be causal (5 are in the >95% list), that are not located within known motifs, annotated elements, nor in any experimentally determined ChIPseq peaks or eQTL credible sets yet discovered. While we have identified a statistically compelling set of genuine associations (often intronic or within 10 kb of strong candidate genes), we can make little inference about function. For example, the single variant credible set only 500 bp from the transcription start site of JAK2 has no annotation, eQTL or ChIPseq peak of note. This underscores the incompleteness of our knowledge regarding the function of non-coding DNA and its role in disease. That the majority of the best refined non-coding associations have no available annotation is perhaps sobering with respect to how well we may be able to currently interpret non-coding variation in medical sequencing efforts. It does suggest, however, that detailed fine-mapping of GWAS signals down to single variants, combined with emerging high-throughput genome-editing methodology, may be among the most effective ways to advance to a greater understanding of the biology of the non-coding genome.

**Supplementary Information** is available in the online version of the paper.

## Acknowledgements

M.J.D. and R.J.X. acknowledge grant supports from P30DK43351, U01DK062432, R01DK64869, Helmsley grant 2015PG-IBD001 and CCFA. C.A.A. and J.C.B are supported by Wellcome Trust grant 098051. M.G. acknowledges grant support from WELBIO (CAUSIBD), BELSPO (BeMGI), Fédération Wallonie-Bruxelles (ARC IBD@Ulg), and Région Wallonne (CIBLES, FEDER). H.H. acknowledges the ASHG/Charles J. Epstein Trainee Award. J.L. acknowledges Wellcome Trust grant 098759/Z/12/Z. D.M. is supported by the Olle Engkvist Foundation and Swedish Research Council (grants 2010–2976 and 2013–3862). R.K.W. is supported by a VIDI grant (016.136.308) from the Netherlands Organization for Scientific Research (NWO). J.D.R. holds a Canada Research Chair and this work was supported by grants from the U.S. National Institute of Diabetes and Digestive and Kidney Diseases (DK064869; DK062432), a grant (CIHR #GPG-102170) from the Canadian Institutes of Health Research to the “CIHR Emerging Team in Integrative Biology of Inflammatory Diseases”, and a *Large-Scale Applied Research Project in Genomics and Personalized Health* grant (GPH-129341) co-funded by the Government of Canada through Genome Canada and the Ministère de l’économie, de l’innovation et de l’exportation du Québec through Génome Québec as well as the Canadian Institutes of Health Research and Crohn’s Colitis Canada. J.H.C. is funded by U01 DK62429, U01 DK062422, and the Sanford J. Grossman Charitable Trust. R.H.D. holds the Inflammatory Bowel Disease Genetic Research Chair at the University of Pittsburgh and acknowledges grant support from U01DK062420 and R01CA141743. E.D. benefitted from a Marie-Curie Fellowship and A-S.G a from fellowships from the FNRS and Fonds Léon Fredericq. J.H. is supported by the Örebro University Hospital Research Foundation and the Swedish Research Council (grant no. 521 2011 2764). M.P. acknowledges the NIHR Biomedical Research Centre awards to Guy’s & St Thomas’ NHS Trust / King’s College London and to Addenbrooke’s Hospital / University of Cambridge School of Clinical Medicine. D.E.: this work was supported by the German Federal Ministry of Education and Research (BMBF) within the framework of the e:Med research and funding concept (SysInflame grant 01ZX1306A). This project received infrastructure support from the DFG Excellence Cluster No. 306 “Inflammation at Interfaces”. A.F. receives an endowment professorship by the Foundation for Experimental Medicine (Zuerich, Switzerland). Additional acknowledgements for the original data are in the Supplementary Information.

## Author Contributions

Overall project supervision and management: M.J.D. J.C.B, M.G. Fine-mapping algorithms: H.H., M.F., L.J. TFBS analyses: H.H., K.F. Epigenetic analyses: M.U.M., G.T. eQTL dataset generation: E.L., E.T., J.D., E.D., M.E., R.M., M.M., Y.M., V.D., A.G. eQTL analyses: M.F., J.D., L.J., A.C. Variance component analysis: T.M., M.F. Contribution to overall statistical analyses: G.B. Primary drafting of the manuscript: M.J.D., J.C.B, M.G., H.H., L.J. Major contribution to drafting of the manuscript: M.F., M.U.M., J.H.C., D.P.M., J.D.R., C.G.M., R.H.D., R.K.W. The remaining authors contributed to the study conception, design, genotyping QC and/or writing of the manuscript. All authors saw, had the opportunity to comment on, and approved the final draft.

## Author Information

The authors declare no competing financial interests. Correspondence and requests for materials should be addressed to H.H., M.G., M.J.D. or J.C.B.

## Methods

### Genotyping and QC

We genotyped 35,197 unaffected and 35,346 affected individuals (20,155 Crohn’s disease and 15,191 ulcerative colitis) using the Immunochip array. Genotypes were called using optiCall^37^ for 192,402 autosomal variants before QC. We removed variants with missing data rate >2% across the whole dataset, or >10% in any one batch, and variants that failed (FDR < 10^−5^ in either the whole dataset or at least two batches) tests for: a) Hardy-Weinberg equilibrium; b) differential missingness between cases and controls; c) significant heterogeneity in allele frequency across controls from different batches. We also removed noncoding variants that were not in the 1000 Genomes Phase I integrated variant set (March 2012 release), or the HapMap phase 2 or 3 releases, as these mostly represent false positives included on Immunochip from the 1000 Genomes pilot, which often genotype poorly. Where a variant failed in exactly one batch we set all genotypes to missing for that batch (to be reimputed later) and included the site if it passed in the remainder of the batches. We removed individuals that had >2% missing data, had significantly higher or lower (defined as FDR<0.01) inbreeding coefficient (*F*), or were duplicated or related (PI_HAT ≥ 0.4, calculated from the LD pruned dataset described below), by sequentially removing the individual with the largest number of related samples until no related samples remain. After QC, there were 67,852 European-derived samples with valid diagnosis (healthy control, Crohn’s disease or ulcerative colitis), and 161,681 genotyped variants available for downstream analyses.

### Linkage-disequilibrium pruning and principal components analysis

From the clean dataset we removed variants in long range LD^38^ or with MAF < 0.05, and then pruned 3 times using the ‘--indep’ option in PLINK (with window size of 50, step size of 5 and VIF threshold of 1.25). This pruned dataset (18,123 variants) was used to calculate the relatedness of the individuals and the principal components. Principal component axes were generated within controls using this LD pruned dataset. The axes were then projected to cases to generate the principal components for all samples. The analysis was performed using our in-house C code (https://github.com/hailianghuang/efficientPCA) and LAPACK package^39^ for efficiency.

### Imputation

Imputation was performed separately in each Immunochip high-density region (184 total) from the 1000 Genomes Phase I integrated haplotype reference panel, downloaded from the IMPUTE2 website (Dec 2013 release). We used SHAPEIT (v2.r769)^40,41^ to pre-phase the genotypes, followed by IMPUTE2 (2.3.0)^14,15^ to perform the imputation. There were 388,432 variants having good imputation quality (INFO > 0.4) and were used in the fine-mapping analysis.

### Manual cluster plot inspection

Variants that had posterior probability greater than 50% or in credible sets mapped to ≤ 10 variants were manually inspected using Evoker v2.2^42^. Each variant was inspected by 3 independent reviewers (10 reviewers participated) and scored as pass, fail or maybe. We remove variants that received one or more fails, or received less than 2 passes. 650 out of 905 inspected variants passed this inspection. A further cluster plot inspection flagged two additional failed variants after removing the failed variants from the first inspection and redoing the imputation and analysis.

### Establishing a p-value threshold

We used a multiple testing corrected p-value threshold for associations of 10^−6^, which was established by permutation. We generated 200 permuted datasets by randomly shuffling phenotypes across samples and carried out association analyses for each permutation across all 161,681 variants in our high-density regions. We stored (i) the ensuing 161,681 x 200 point-wise p-values (*α_S_*), as well as (ii) the 200 “best” p-values (*α_B_*) of each permuted datasets. We then computed the empirical, family-wise p-value (*α_M_*)(corrected for multiple testing) for each of the 161,681 x 200 tests as its rank/200 with respect to the 200 *α_B_*. We then estimated the number of independent tests performed in the studied regions, *n*, as the slope of the regression of log(1-*α_M_*) on log(1-*α_S_*), knowing that *α_M_* = 1 − (1 − *α_S_*)*^n^*.

### Detecting and fine-mapping association signals

We used three fine-mapping methods to detect independent signals and create credible sets across 103 high-density regions (Supplementary Methods). Signals identified by different methods were merged if their credible sets shared one or more variants. In order to adjudicate differences between methods, we first assigned each candidate signal to the combination of a lead variant and trait (CD, UC or IBD) that maximizes the marginal likelihood from equation (8) in Supplementary Methods. At loci with >1 signal, we fixed the signals reported by all three methods, and then tested all possible combinations of signals reported by one or two methods, selecting whichever combination has the highest joint marginal likelihood. We consider signals to be confidently fine-mapped, and take them forward for subsequent analysis, if they a) are in loci where the lead variant has p < 10^−6^, b) have a ratio of Bayes factors for the best model and the second best model greater than 10, c) are reported by more than one method and d) passed cluster plot inspection.

### Phenotype assignment of signals

We assign each signal as CD-specific, UC-specific or shared, using the Bayesian multinomial model used for fine-mapping method 2 (the method best able to assess evidence of sharing in the presence of potentially correlated effect sizes). For the lead variant for each credible set, we calculate the marginal likelihoods as in equation 13 from Supplementary Methods, restricting either β_UC_ = 0 (for the CD-only model) or β_CD_ = 0 (for the UC-only model), as well as using the unconstrained prior (for the associated to both model), and select the model with the highest marginal likelihood. We then calculate the log Bayes factor in favor of sharing, i.e. the log of ratio of marginal likelihoods between the associated-to-both model and the best of the single-phenotype associated models.

### Estimating the variance explained by the fine-mapping

We used a mixed model framework to estimate the total risk variance attributable to the IBD risk loci, and to the signals identified in the fine-mapping. We the GCTA software package^43^ to compute a gametic relationship matrix (G-matrix) using genotype dosage information for the genotyped variants in the high-density regions (which we will call **G***_HD_*). We then fit a variety of variance component models by restricted maximum likelihood analysis using an underlying liability threshold model implemented with the DMU package^44^. The first model is a standard heritability mixed-model that includes fixed effects for five principal components (to correct for stratification) and a random effect summarizing the contribution of all variants in the fine-mapping regions, such that the liabilities across all individuals are distributed according to

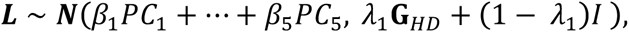

where *λ*_1_ is thus the variance explained by all variants in fine-mapping regions, which we estimate. We then fitted a model that included an additional random effect for the contribution of the lead variants have been specifically identified (with G-matrix **G***_Signals_*), such that liability is distributed

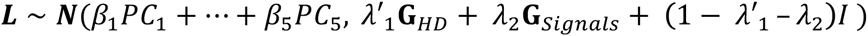

The variance explained by the signals under consideration is then given by the reduction in the variance explained by all variants in the fine-mapping regions between the two models (*λ*_1_ − *λ*′_1_). We used this approach to estimated what fraction of this variance was accounted for by (i) the single strongest signals in each region (as would be typically done prior to fine-mapping), or (ii) the all signals identified in fine-mapping. We used Cox and Snell’s method^45^ to estimate the variance explained across independent signals (Extended Data Figure 2b) for computational efficiency.

### Overlap between transcription factor binding motifs and causal variants

For each motif in the ENCODE TF ChIP-seq data (http://compbio.mit.edu/encode-motifs/, accessed Nov 2014) ^20^, we calculated the overall information content (IC) as the sum of IC for each position^46^, and only considered motifs with overall IC ≥ 14 bits (equivalent to 7 perfectly conserved positions). For every variant in a high-density region we determined if it creates or disrupts a motif at a high-information site (IC ≥ 1.8). For each credible set that contains a motif-affecting variant, we calculated a p-value as 1 − (1 − *f*)*^n^*, where *n* is the size of the credible set and *f* is the proportion of all variants in the high-density region that disrupt or a create a motif in that TF family.

### Overlap between epigenetic signatures and causal variants

For each combination of 120 tissues and three histone marks (H3K4me1, H3K4me3 and H3K27ac) from the Roadmap Epigenome Project we calculated an overlap score, equal to the sum of fine-mapping posterior probabilities for all variants in peaks of that histone mark in that tissue. We generated a null distribution of this score for each tissue/mark by shifting chromatin marks randomly over the high-density regions (shifting the peaks from their actual position by a random number of bases while keeping inter-peak spacing the same) and calculating the overlap score for each permutation. To summarize these correlated results across many cell and tissue types we defined a set of “core” H3K4me1 immune and H3K27ac gut peaks as sets of overlapping peaks in cells that showed the strongest enrichment (p<10^−4^). Intersects were made using bedtools v2.24.0 default settings^47^. We selected 6 immune cell types for H3K4me1 and 3 gut cell types for H3K27ac (Supplementary Table 2). We also chose controls (Supplementary Table 2) from non-immune and non-gut cell types with similar density of peaks in the fine-mapped regions as compared to immune/gut cell types to confirm the tissue-specificity of the overlap. We used the phenotype assignments (described above) in dissecting the enrichment for the CD and UC signals. Sixty-five CD and 21 UC signals were used in this analysis.

### Published eQTL summary statistics

We used eQTL summary statistics from two published studies:

- Peripheral blood eQTLs from the GODOT study^48^ of 2,752 twins, reporting loci with MAF>0.5%.
- CD14+ monocyte eQTLs from Table S2 in Fairfax *et al*.^31^, comprised of 432 European individuals, measured in a naïve state and after stimulation with interferon-*γ* (for 2 or 24 hours) or lipopolysaccharide. Reports loci with MAF>4% and FDR<0.05.

### Processing and quality control of new eQTL ULg dataset

A detailed description of the ULg dataset is in preparation (Momozawa et al., in preparation). Briefly, we collected venous blood and intestinal biopsies at three locations (ileum, transverse colon and rectum) from 350 healthy individuals of European descent, average age 54 (range 17–87), 56% female. SNPs were genotyped on Illumina Human OmniExpress v1.0 arrays interrogating 730,525 variants, and SNPs and individuals were subject to standard QC procedures using call rate, Hardy-Weinberg equilibrium, MAF ≥ 0.05, and consistency between declared and genotype-based sex as criteria. We further imputed genotypes at ~7 million variants on the entire cohort using the Impute2 software package^14^ and the 1,000 Genomes Project as reference population (Phase 3 integrated variant set, released 12 Oct 2014) ^11,15^. From the blood, we purified CD4+, CD8+, CD19+, CD14+ and CD15+ cells by positive selection, and platelets (CD45-negative) by negative selection. RNA from all leucocyte samples and intestinal biopsies was hybridized on Illumina Human HT-12 arrays v4. After standard QC, raw fluorescent intensities were variance stabilized^49^ and quantile normalized^50^ using the lumi R package^51^, and were corrected for sex, age, smoking status, number of probes with expression level significantly above background as fixed effects and array number (sentrix id) as random effect. For each probe with measureable expression (detection p-value < 0.05 in >25% of samples) we tested for cis-eQTLs at all variants within a 500 kilobase window. The nominal p-value of the best SNP within a cis-window was Sidak-corrected for the window-specific number of independent tests, and we estimated false discovery rates (q-values) from the resulting p-values across all probes using the qvalue R package^52^. 480 cis-eQTL with FDR ≤ 0.10 with the lead SNPs within the 97 high-density regions (94 fine-mapped plus 3 unresolved) were retained for further analyses.

### Naïve co-localization using lead SNPs

We calculated the proportion of IBD credible sets that contain a lead eQTL variant in a particular tissue. This value is then compared to a background rate:

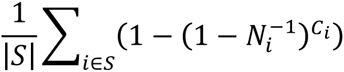

where *N_i_* is the total number of variants in region *i* in 1000 Genomes with an allele frequency greater than a certain threshold (equal to the threshold used for the original eQTL study), *C_i_* is the number of these variants that lie in IBD credible sets, and *S* is a set of regions that have at least one significant eQTL (|*S*| is the number of regions in this set). P-values can then be calculated assuming a binomial distribution with probability equal to the background rate and the number of trials equal to |*S*|.

### Frequentist co-localization using conditional p-values

We next used conditional association to test for evidence of co-localization, as described in Nica et al.^25^. This method compares the p-value of association for the lead SNP of an eQTL before and after conditioning on the SNP with the highest posterior in the credible set, and measures the drop in log(1/p). An empirical p-value for this drop is then calculated by comparing it to the drop for all variants (with MAF ≥ 0.05) in the high-density region. An empirical p-value ≤ 0.05 was considered as evidence that the corresponding credible set is co-localized with the corresponding cis-eQTL. To evaluate whether our 139 credible sets affected cis-eQTL more often than expected by chance we counted the number of credible sets affecting at least one cis-eQTL with p-value ≤ 0.05, and compared how often this number was matched or exceeded by 1,000 sets of 139 lead variants that were randomly selected yet distributed amongst the 94 loci in accordance with the real credible sets.

### Bayesian co-localization using Bayes factors

Finally, we used the Bayesian co-localization methodology described by Giambartolomei et al^53^, modified to use the credible sets and posteriors generated by our fine-mapping methods. The method takes as input a pair of IBD and eQTL signals, with corresponding credible sets *S^IBD^* and *S^eQTL^*, and posteriors for each variant 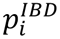 and 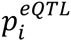 (with 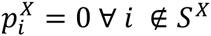). Credible sets and posteriors were generated for eQTL signals using the Bayesian quantitative association mode in SNPTest (with default parameters), with credible sets in regions with multiple independent signals generated conditional on all other signals. Our method calculates a Bayes factor summarizing the evidence in favor of a colocalized model (i.e. a single underlying causal variant between the IBD and eQTL signals) compared to a non-colocalized model (where different causal variants are driving the two signals), given by the ratio of marginal likelihoods

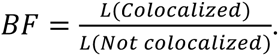

The marginal likelihood for the colocalized model (i.e. hypothesis *H*_4_ in Giambartolomei et al) is given by

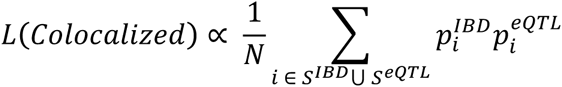

and the likelihood for the model where the signals are not colocalized (i.e., hypothesis *H*_3_) is given by:

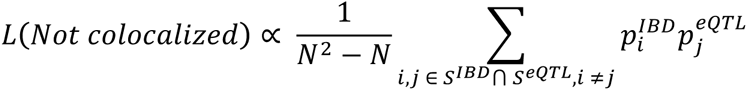

In both cases, N is the total number of variants in the region. We only count towards N variants that have *r*^2^ > 0.2 with either the lead eQTL variant or the lead IBD variant.

*Permutation analysis.* To measure enrichment in colocalization Bayes factors compared to the null, we carried out a permutation analysis. In this analysis, we randomly reassigned eQTL signals to new fine-mapping regions to generate a set of simulated null datasets. This is carried out using the following scheme:

1. Estimate the standarized effect size *β_g_* for each eQTL signal *g*, equal to standard deviation increase in gene expression for each dose of the minor allele.
2. Randomly reassign each eQTL signal to a new fine-mapping region, and then select a new causal variant with a minor allele frequency within 1 percentage point of the lead variant from the real signal. If multiple such variants exist, select one at random. If no such variants exist, pick the variant with the closest minor allele frequency.
3. Generate new simulated gene expression signals for each individual from 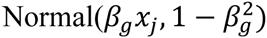 where *x_j_* is the individual’s minor allele dosage at the new causal variant and f is the minor allele frequency.
4. Carry out fine-mapping and calculate colocalization Bayes factors for each pair of (real) IBD signal and (simulated) eQTL signal.
5. Repeat stages 2–4 1000 times for each tissue type

We can use these permuted Bayes factors to calculate p-values for each IBD credible set, given by the proportion of time the permuted BFs were as large or greater than the one observed in the real dataset. To generate a high-quality set of colocalized eQTL and IBD signals, we take all signals that have BF > 2, p < 0.01 and *r*^2^ between hits of >0.8.

---

**Extended Data Figure 1.**
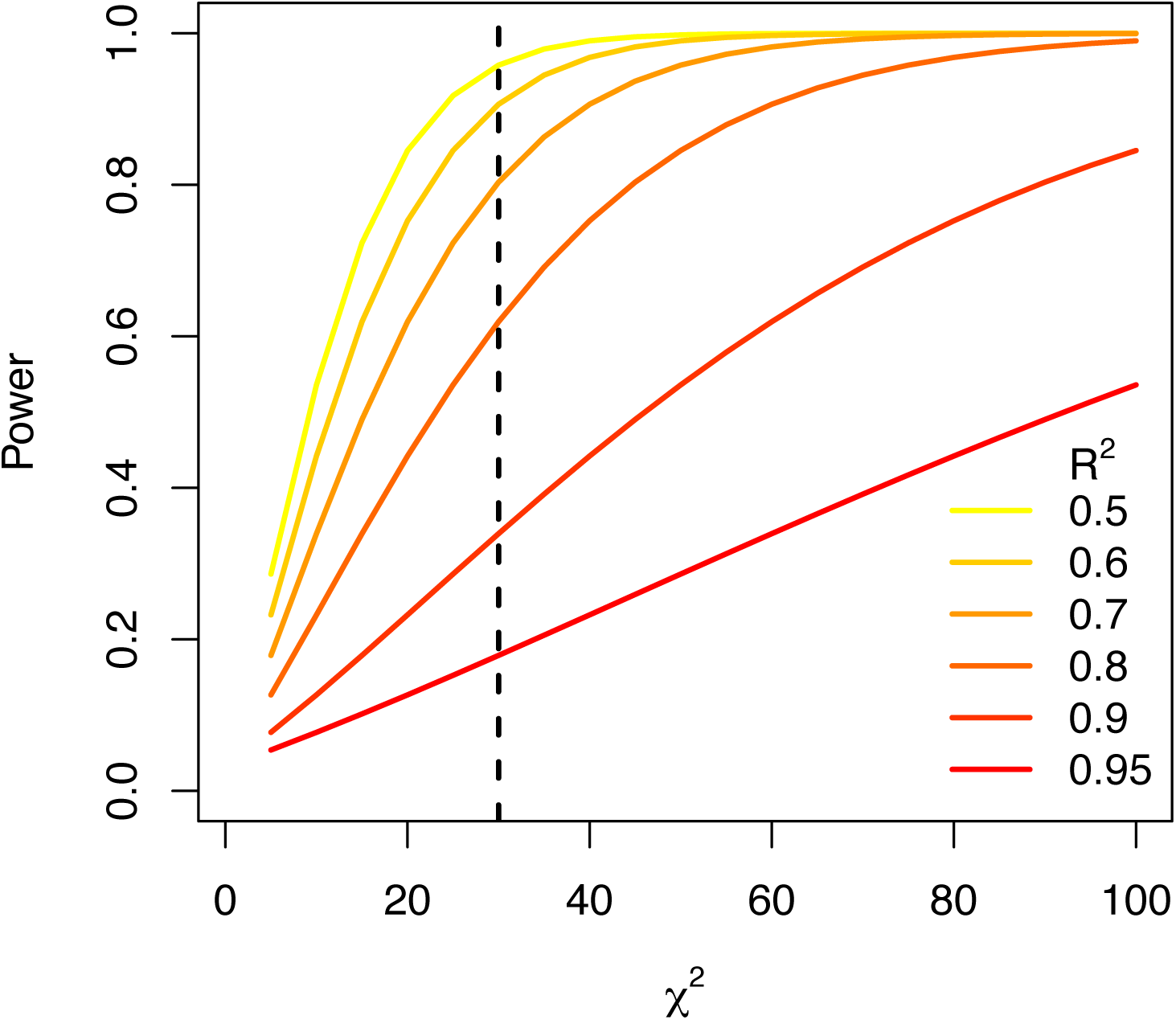
Power (y axis) to distinguish which variant in a correlated pair (strength of correlation shown by color) is causal increases with the significance of the association (x axis), and therefore with sample size and effect size. The vertical dashed line flags the genome-wide significance level. To estimate the relationship between the strength of association and our ability to fine-map it, we assumed that the association has only two possible causal variants, and we define the signal as successfully fine-mapped if the ratio of Bayes factors between the true causal variant and the non-causal variant is greater than 10 (a 91% posterior, assuming equal priors). Using equation (8) in Supplementary Methods, we have

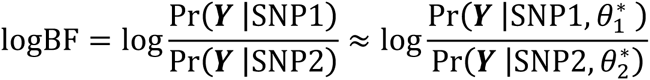

in which *θ*^*^ is maximum likelihood estimate of the parameter values. The log-likelihood ratio follows a chi-square distribution:

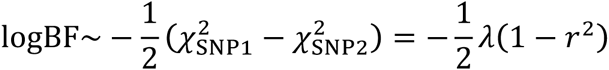

in which *λ* is the chi-square statistic of the lead variant and *r* is the correlation coefficient between the two variants. Because of the additive property of the chi-square distribution, logBF follows a non-central chi-square distribution with 1 degree of freedom and non-centrality parameter *λ*(1 − *r*^2^)/2. Therefore, the power can calculated as the probability that logBF > log(10), given by the CDF of the non-central chi-squared distribution.

**Extended Data Figure 2.**
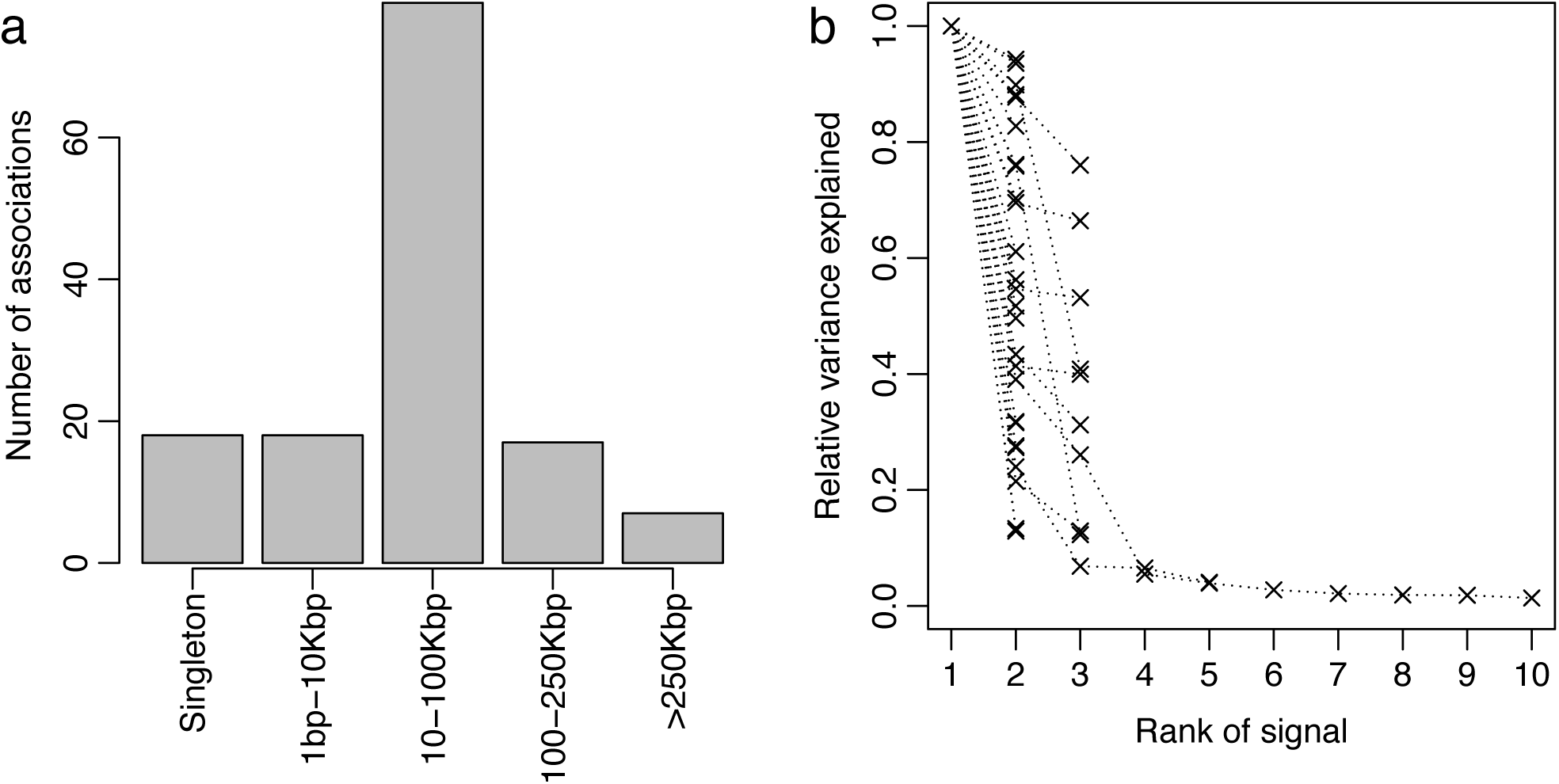
**a**, Genomic distance that variants in 95% credible set span. **b**, Variance explained normalized to the primary association in each locus.

**Extended Data Figure 3.**
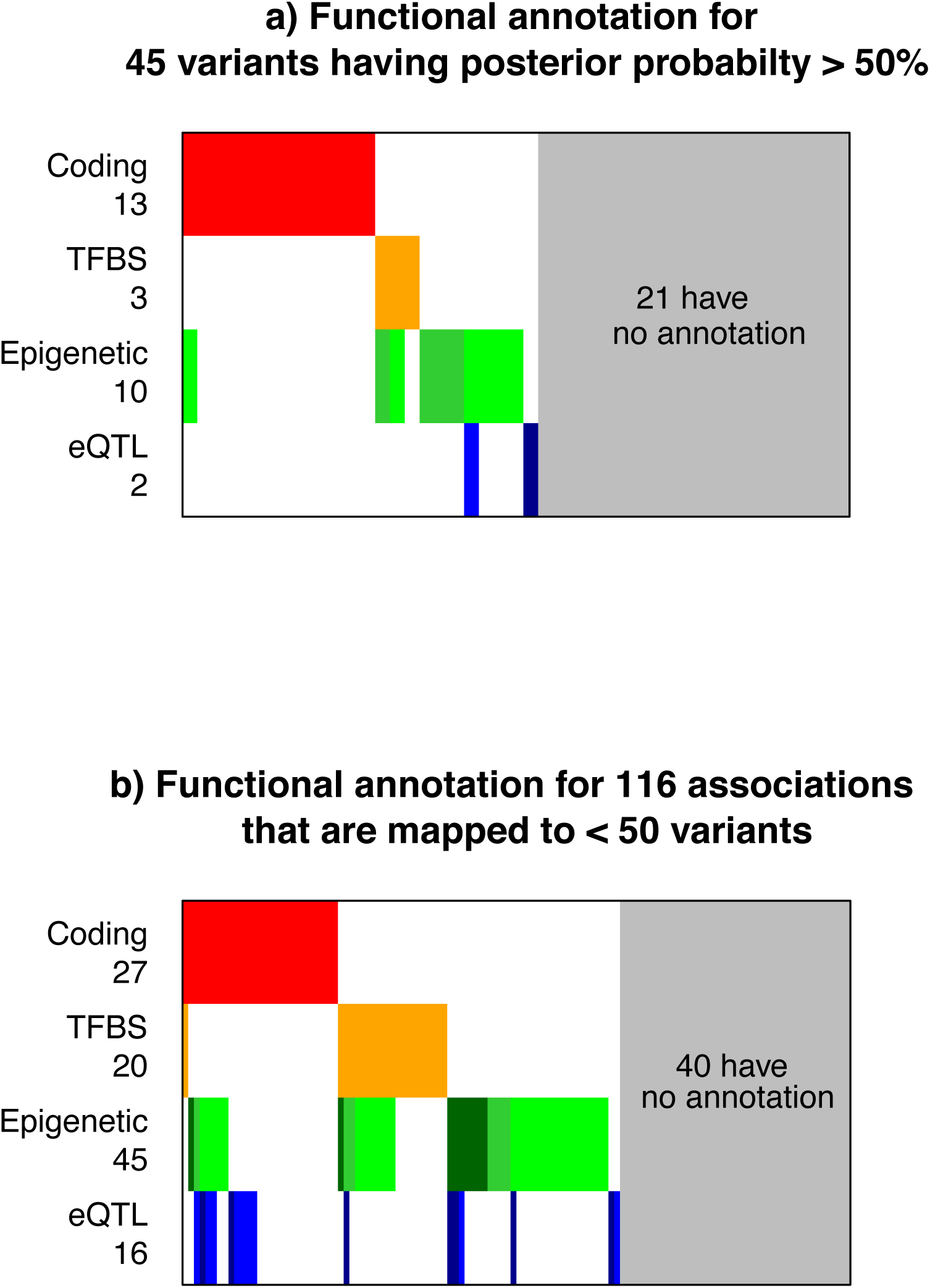
**a**, Functional annotation for 45 variants having posterior probability > 50%. **b**, Functional annotation for 116 associations that are fine-mapped to ≤ 50 variants.

**Extended Data Figure 4.**
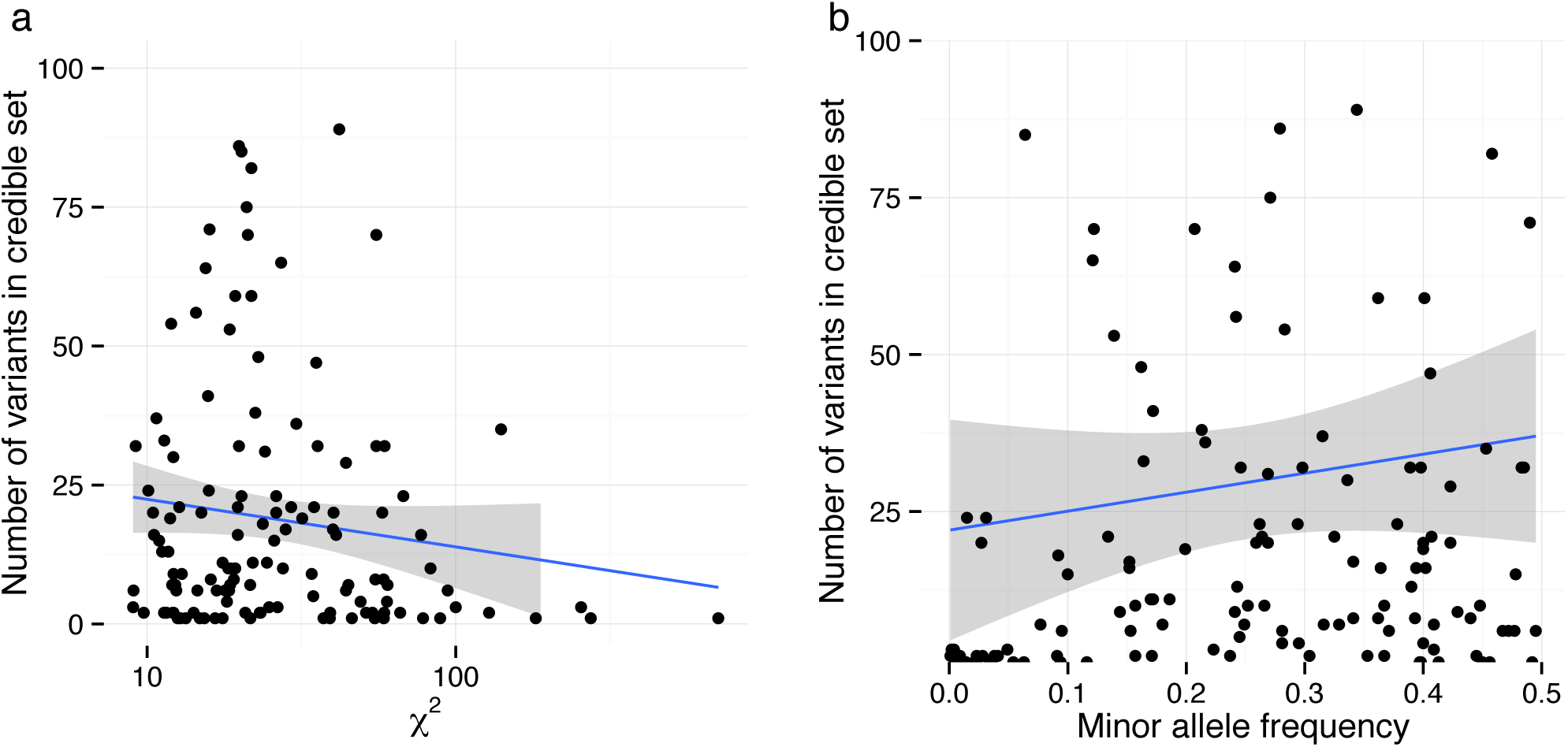
**a**, Number of variants in credible set decreases with the significance of the signal. **b**, Number of variants in credible set increases with the minor allele frequency of the signal. The solid line shows the fitted trend in both panels, and the shaded region shows the variance of the trend.

**Extended Data Figure 5.**
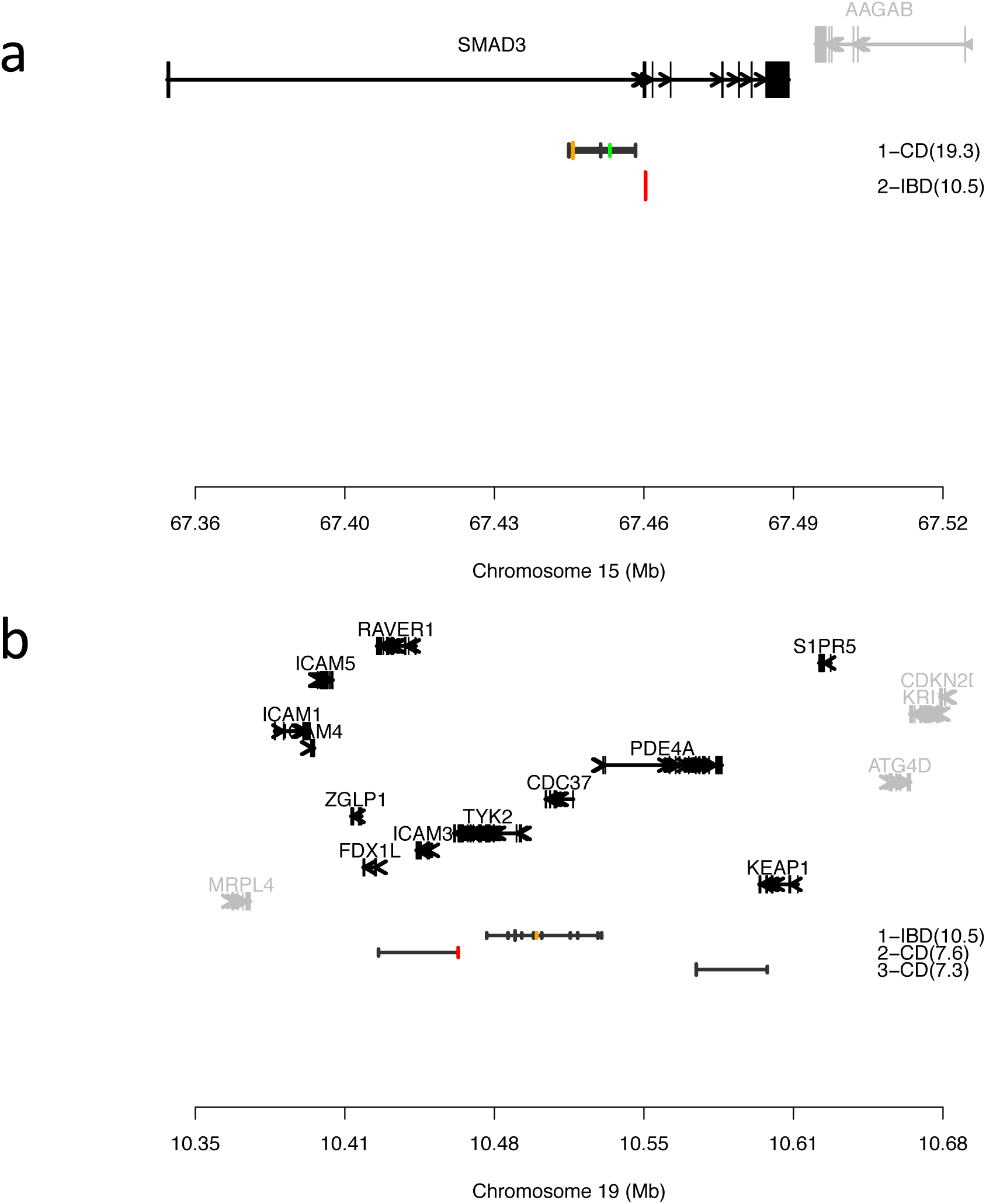
SMAD3 (**a**) and TYK2 (**b**) regions after fine-mapping. The implicated region has been reduced to a smaller number of genes (shown in black). Color ticks are variants mapped to their functions (red: non-synonymous, orange: disrupting TFBS, green: overlapping epigenetic marks) and black ticks are variants not mapped to a function. The width of the tick scales with the posterior probability.

**Extended Data Figure 6.**
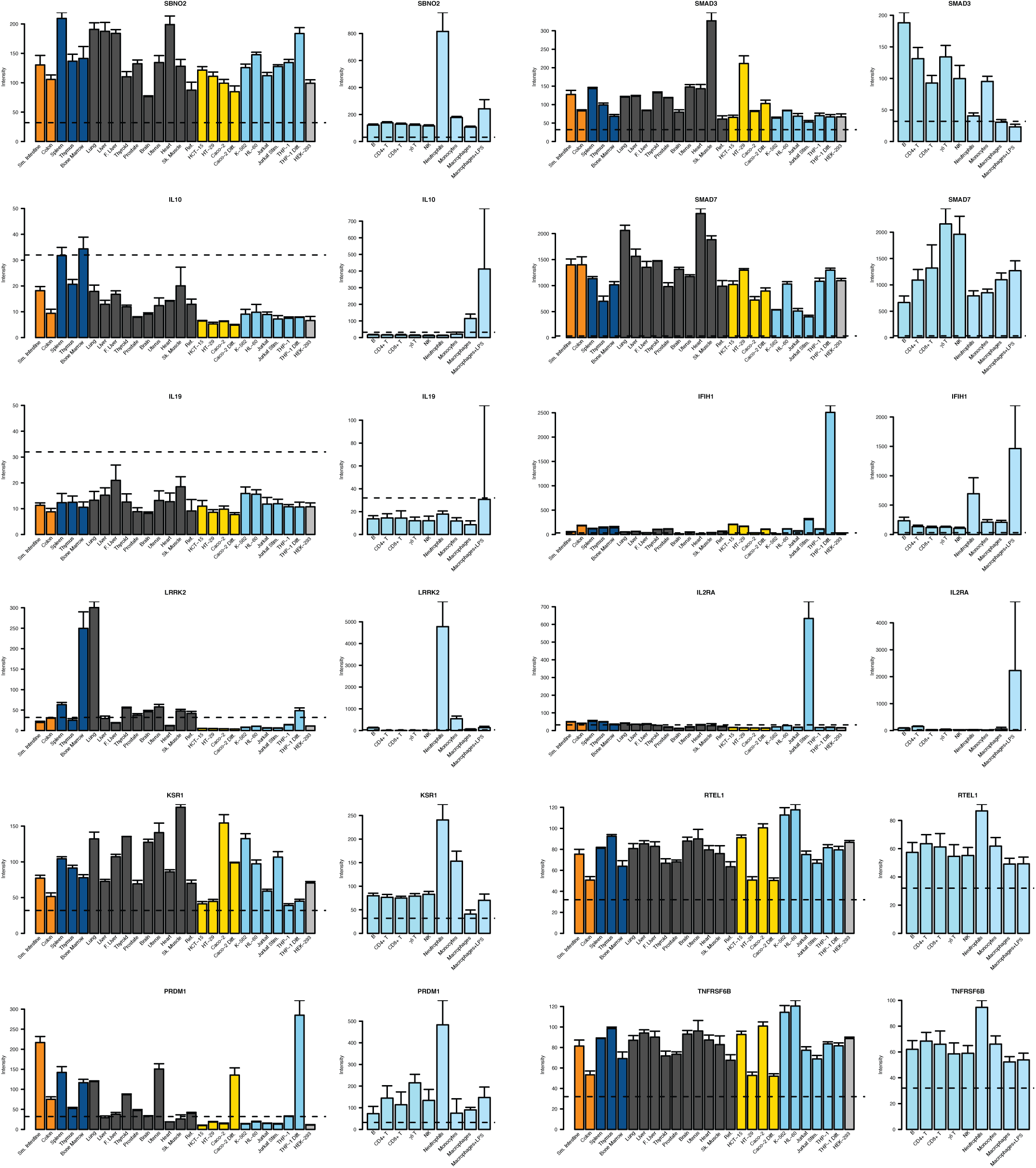
Tissue and cell line specific expression for genes *SBNO2, IL10, IL19, LRRK2, KSR1, PRDM1, SMAD3, SMAD7, IFIH1*, *IL2RA, RETL1* and *TNFRSF6B.* **Left panels.** Expression levels of selected genes were determined in a panel of human tissues (bone marrow, heart, skeletal muscle (Sk. Muscle), uterus, liver, fetal liver (F. Liver), spleen, thymus, thyroid, prostate, brain, lung, small intestine (Sm. Intestine) and colon) and human cell lines using a custom made Agilent expression array. The cell lines represent models of human T lymphocytes (Jurkat), monocytes (THP-1), erythroleukemia cells (K562), promyelocytic cells (HL-60), colonic epithelial cells (HCT-15, HT-29, Caco-2), and cells from embryonic kidney (HEK-293). In addition, models of differentiated colonic epithelium (Caco-2 differentiated for 21 days in culture (Caco-2 diff.)), activated T lymphocytes (Jurkat cells stimulated with PMA (40ng/ml) and ionomycin (1ug/ml) for 6 hrs (Jurkat stim.)), and macrophages (derived from THP-1 differentiated for 24 hrs (THP-1 diff.) with IFN-γ (400U/ml) and TNF-α (10ng/ml)) were examined. Intensity values for each tissue/cell line represent the geometric mean with geometric standard deviation of 3 independent measurements; each measurement represents the geometric mean of all probes (one per exon) for each gene followed by a median normalization across all genes on the array. The dotted line indicates the threshold level for detection of basal expression. The reference sample (Ref.) is composed of a mixture RNAs derived from 10 different human tissues. **Right panels.** Expression levels of selected genes were determined in a panel of primary immune cells (neutrophils, monocytes, γδ T cells, B cells, NK cells, CD4^+^ T cells, CD8^+^ T cells) isolated from healthy donors, as well as monocyte *in vitro* derived macrophages without and with 24 hours of stimulation using 1 ug/ml of lipopolysaccharide (macrophages+LPS). The results presented in the **left and right panels** were generated and analyzed separately and therefore the expression values are not directly comparable.

**Extended Data Table 1.**
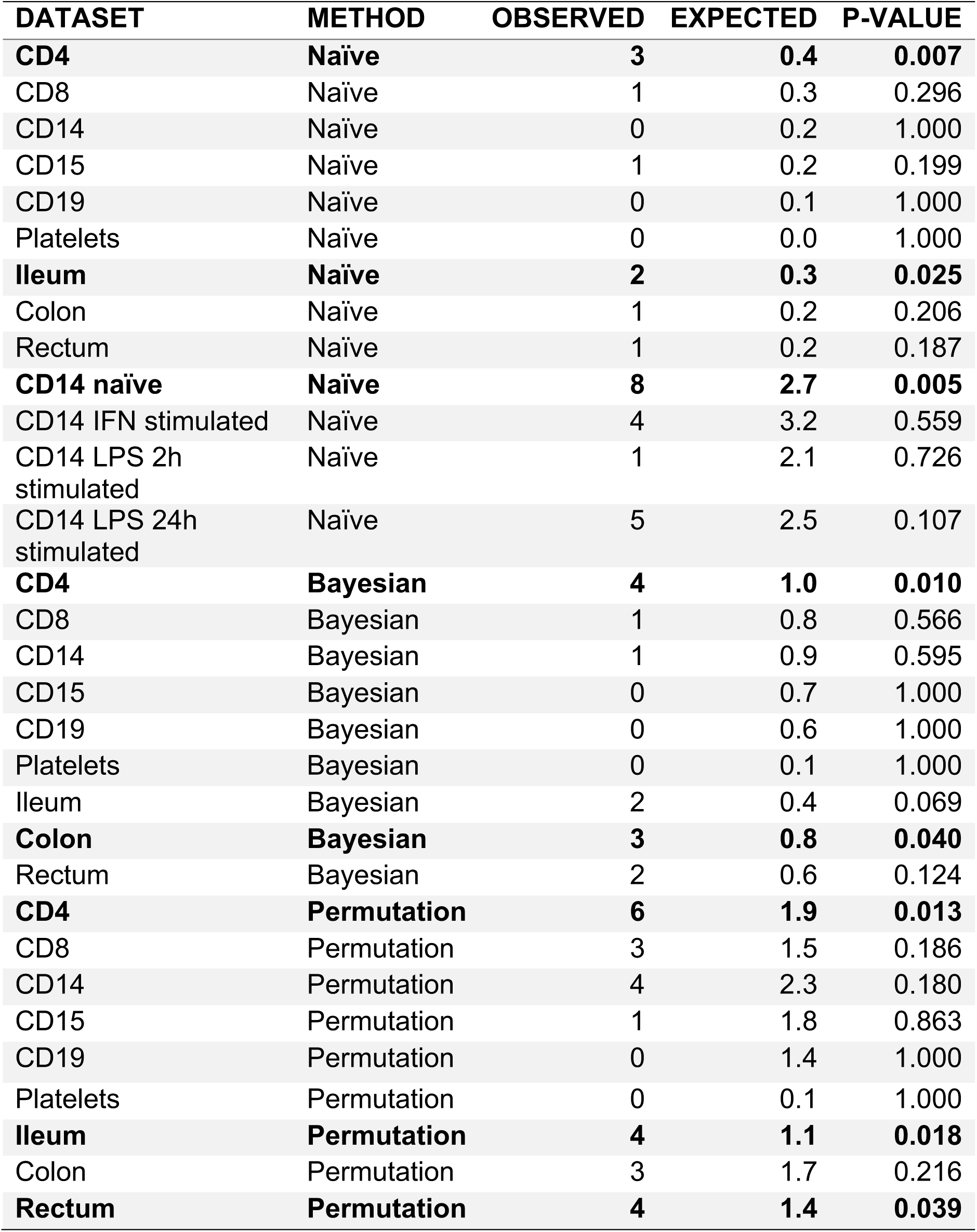
The number IBD credible sets that colocalize with expression QTLs using the naïve, permutation-based and Bayesian-based approaches.

